# Hair cells with heterogeneous transfer characteristics encode mechanical stimuli in the lateral line of zebrafish

**DOI:** 10.1101/261669

**Authors:** Paul Pichler, Leon Lagnado

## Abstract

Ribbon synapses of hair cells transmit mechanical information but the transfer characteristics relating deflection of the hair bundle to glutamate release have not been assessed directly. Here we have imaged glutamate to investigate how hair cells encode information in the lateral line of zebrafish. Half the hair cells signalled cupula motion in either direction from rest, achieving maximum sensitivity for deflections of ~40 nm in the preferred direction. The remainder rectified completely and were less sensitive, extending the operating range of the neuromast beyond 1μm. Adaptation was also heterogeneous, with some hair cells generating sustained synaptic outputs and others transient. A unique signal encoded a return to rest: a transient burst from hair cells unresponsive to the initial stimulus. A mixed population of hair cells with these various transfer characteristics will allow a neuromast to encode weak stimuli as well as the amplitude and duration of stronger deflections.

## Introduction

An increasingly important context for the study of mechanotransduction is the lateral line of larval zebrafish (Graydon et al 2017; Oteiza et al 2017; Sheets et al 2017; Erickson et al 2017; Maeda et al 2017). This sensory system enables the animal to detect vibrations and pressure gradients in its hydrodynamic environment and is implicated in social behaviours (Butler & Maruska 2016), predator avoidance (McHenry et al 2009; Stewart et al 2013) and the maintenance of position against a current (rheotaxis; Olive et al 2016; Oteiza et al 2017). The sense organs of the lateral line, called neuromasts, are distributed over the head and body of the zebrafish, each being composed of 10-20 hair cells (Pujol-Martí & López-Schier 2013). As in the auditory and vestibular systems, the hair cells convert small displacements of the hair bundle into changes in the rate of glutamate release from ribbon-type synapses (Nicolson, 2015). Understanding the transfer characteristics of these synapses is therefore fundamental to understanding how mechanical information is encoded within the lateral line.

All the hair cells within a neuromast act as a population to encode a single mechanical stimulus – the deflection of the cupula into which they all project. The direction of deflection is encoded by a “push-pull” system in which half the hair cells are depolarized by motion in one direction and the other half by motion in the opposite, with segregation of these two populations onto separate afferent fibers (Faucherre et al 2009). A number of fundamental questions about signalling in the lateral line remain unanswered. How does the output of individual hair cells encode deflections of the cupula? What is the dynamic range over which signalling occurs? And how does the output from the synaptic ribbon adapt? It is equally important to understand how these properties might vary between hair cells to determine how the population acts to encode the amplitude and duration of a stimulus.

Direct answers to these questions require assaying the release of glutamate from individual hair cells while deflecting the hair bundle by known amounts. The output of hair cells has been studied by measuring capacitance changes (Beutner et al 2001; Brandt et al 2005; Ricci et al 2013; Olt et al 2014) or by recording synaptic currents in the afferent fiber (Keen & Hudspeth 2006; Li et al 2009; Weisz et al 2012), but both these techniques require the synapse to be activated by direct injection of current which bypasses the normal process of mechanotransduction. We therefore developed an all-optical approach in which deflections of the cupula and release of glutamate from multiple hair cells were imaged within one neuromast. Monitoring synaptic transmission using the glutamate sensor iGluSnFR (Marvin et al 2013) allowed us to investigate the transfer characteristics of both individual hair cells and the population.

Here we show that the output of a neuromast is determined by a heterogeneous population of hair cells. About 40% signal deflections of the hair bundle less than 100 nm, while the remainder are less sensitive and extend the dynamic range of the neuromast to encode deflections beyond 1 μm. These heterogeneous transfer characteristics extend to the dynamics of adaptation, enabling the generating of a population code that signals weak stimuli while maintaining the ability to encode the amplitude and duration of stronger deflections.

## Results

### An all-optical approach to measuring the transfer characteristics of hair cell ribbon synapses *in vivo*

The lateral line system has been studied intensively but the input-output relation of hair cells within neuromasts is still unclear. To observe the output in larval zebrafish we used the *Sill* promoter to drive expression of the fluorescent glutamate sensor iGluSnFR in the surface membrane of primary afferents postsynaptic to hair cell ribbons (Pujol-Martí et al 2012; Marvin et al 2013; Figs. 1A-C). Neuromasts in the posterior lateral line were stimulated using a narrow pipette that applied positive and negative pressure steps, generating iGluSnFR signals at distinct hotspots (Fig. 1D-F). To test if these hotspots coincided with presynaptic ribbons we used fish co-expressing iGluSnFr with Ribeye-mCherry (Odermatt et al 2012). Figs. 1D and E home in on four varicosities from two afferents: two of the ROIs were contacted by one ribbon (amber and blue ROIs) while the other two were not (red and green ROI). iGluSnFR signals were only observed in areas that were not in close apposition to a ribbon synapse (Fig. 1F), and this was the rule in all three neuromasts in which this test was made. This example also demonstrates how synapses of opposite polarity could be monitored simultaneously: while the blue ROI was activated by positive deflections towards the head the amber ROI was activated by negative deflections towards the tail (Fig. 1F).

**Figure 1.**
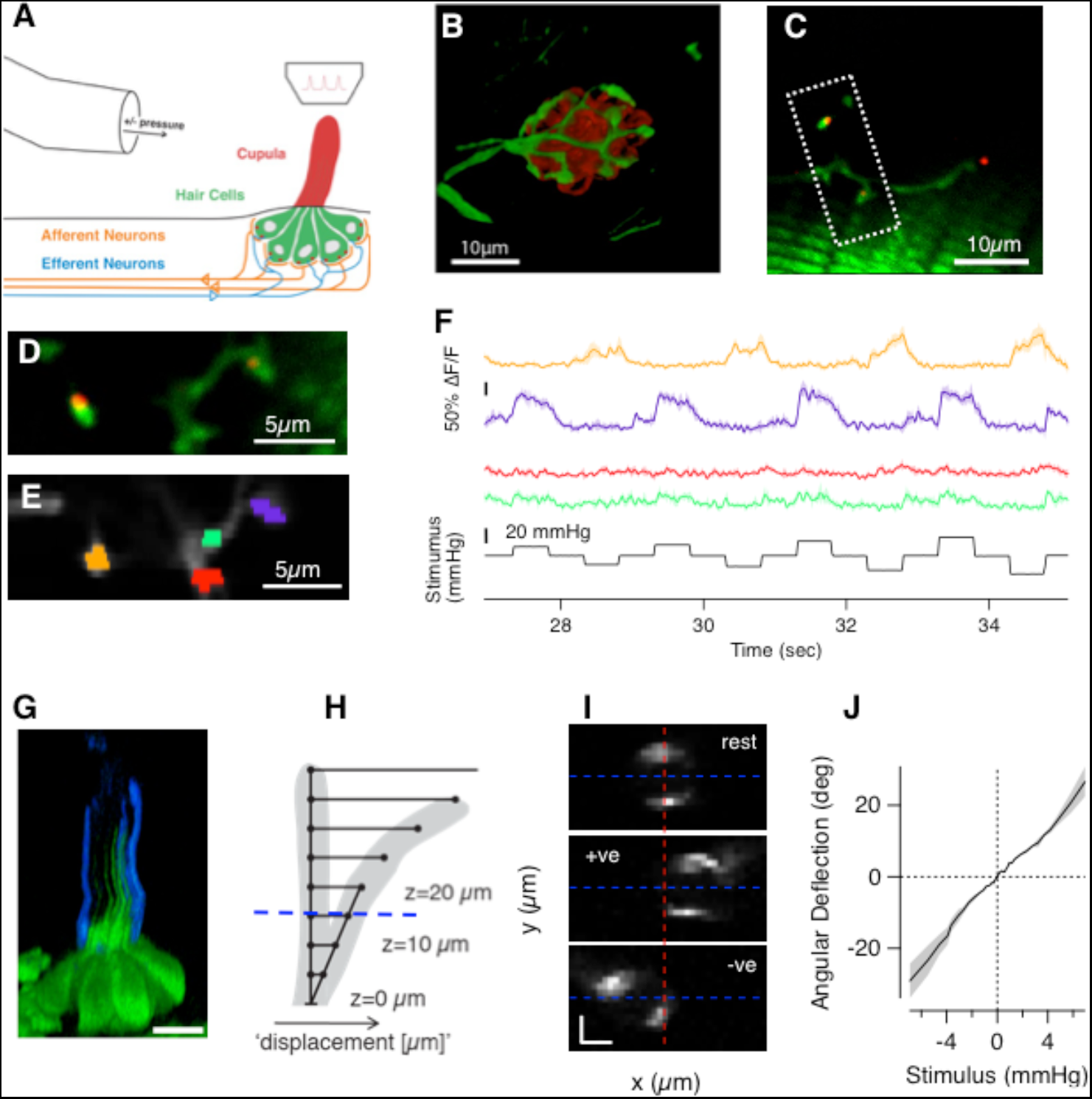
Imaging the input and output from hair cells in a neuromast. (**A**) The NM was stimulated with positive and negative pressure steps applied through a bent pipette while being imaged through a two-photon microscope. (**B**) iGluSnFR was expressed over the surface of afferent neurons (green) which form a basket like structure around hair cells, here counterstained in red using FM4-64. (**C**) Image of a NM in a 7dpf larva (Tg[Sill2, UAS::iGluSnFR, Rib::Rib-mCherry]) showing varicosities of an afferent neuron (green) as well as synaptic ribbons (red). (**D**) Close-up of the boxed area in C. Of the four visible varicosities, two did not coincide with presynaptic ribbons. (**E**) Regions of interest (ROIs) used for analysis. (**F**) The responses of the four ROIs in E. The amber and blue ROIs postsynaptic to ribbons responded to positive and negative pressure steps, respectively. The red and green ROIs not apposed to ribbons did not respond. (**G**) Side view of a NM of expressing GFP in hair cells with the surface of the cupula stained with Alexa-350-WGA. Notice how the kinocilia extend roughly half way up the cupula (scale bar 10 μm). (**H**) Schematic of the model used to calculate angular deflections of the cupula from a pivot at its base. The translational deflection for a given stimulus pressure was measured at several distances z above the apical surface of the hair cell. (**I**) Three representative frames of the stained cupula at z = 15 μm at rest and deflected by a positive and negative pressure step. The angular deflection was calculated as tan^-1^ (Δx/z), where Δx was the translation in the centre of mass of the staining. **(J)** Example of a calibration curve relating the stimulus pressure to the angular deflection. These relations were generally linear.

To quantify the mechanical input to the hair cells, deflections of the cupula were visualized by staining polysaccharides on the surface blue or red with Alexa-350/594 coupled Wheat Germ Agglutinin (Fig. 1G-J; detailed in Experimental Procedures and Supplementary Figure 1). The rotational deflection of the cupula was calculated based on the idea that the proximal region, containing the kinocilia, acts as a rigid lever pivoting on a plane at the apical surface of the hair cells (Fig. 1I; McHenry & van Netten 2007). We tested this model by imposing a variety of pressure steps and tracking the translational motion of the cupula through planes at four different distances from the apical surface of the hair cell (Supplementary Figure 1E). Fig. 1I shows images of the cupula at *z* = 15 μm, from which the *x* translation was estimated from the movement of the centre of mass of the fluorescence. At any single pressure, the x translations at different z gave consistent estimates of the angle of rotation confirming that the lower part of the cupula behaves as a beam pivoting around its base (Supplementary Fig. 1D). We were therefore able to calibrate the relation between applied pressure and rotation of the cupula for each experiment, which in turn allowed us to control for variables such as the diameter of the pipette delivering the stimulus or its location relative to the neuromast (Fig. 1J & Supplementary Figure 1F).

### Ribbon synapses in the lateral line can signal deflections less than 100 nm

What is the mechanical sensitivity of hair cells in the lateral line? Relating spikes in the afferent nerve to estimates of cupula deflection, Haehnel-Taguchi et al (2014) report that deflections below 8 μm cannot be encoded. In contrast, calcium imaging in the hair cells themselves indicate that robust responses can be elicited by deflections of about 2-3 μm (Sheets et al 2012; Kindt et al 2012). Imaging glutamate release we found that many hair cells were at least an order of magnitude more sensitive than these estimates, signalling deflections less than 100 nm.

The traces in Fig. 2A show iGluSnFR signals from two nearby hair cells in response to positive and negative pressure steps of increasing amplitude, each lasting 1 s. It can be immediately seen that hair cell 1 (black trace) was more sensitive to small deflections from rest, generating changes in glutamate release at stimulus strengths that did not elicit glutamate release from hair cell 2 (red trace). The mechanical tuning of these receptors was characterized as the peak amplitude of the iGluSnFR signal (R) as a function of angular rotation of the cupula (X), as plotted in Fig. 2B. A good description of this relation was provided by a two-state Boltzmann equation of the form:

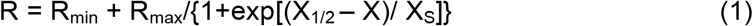

**Figure 2.**
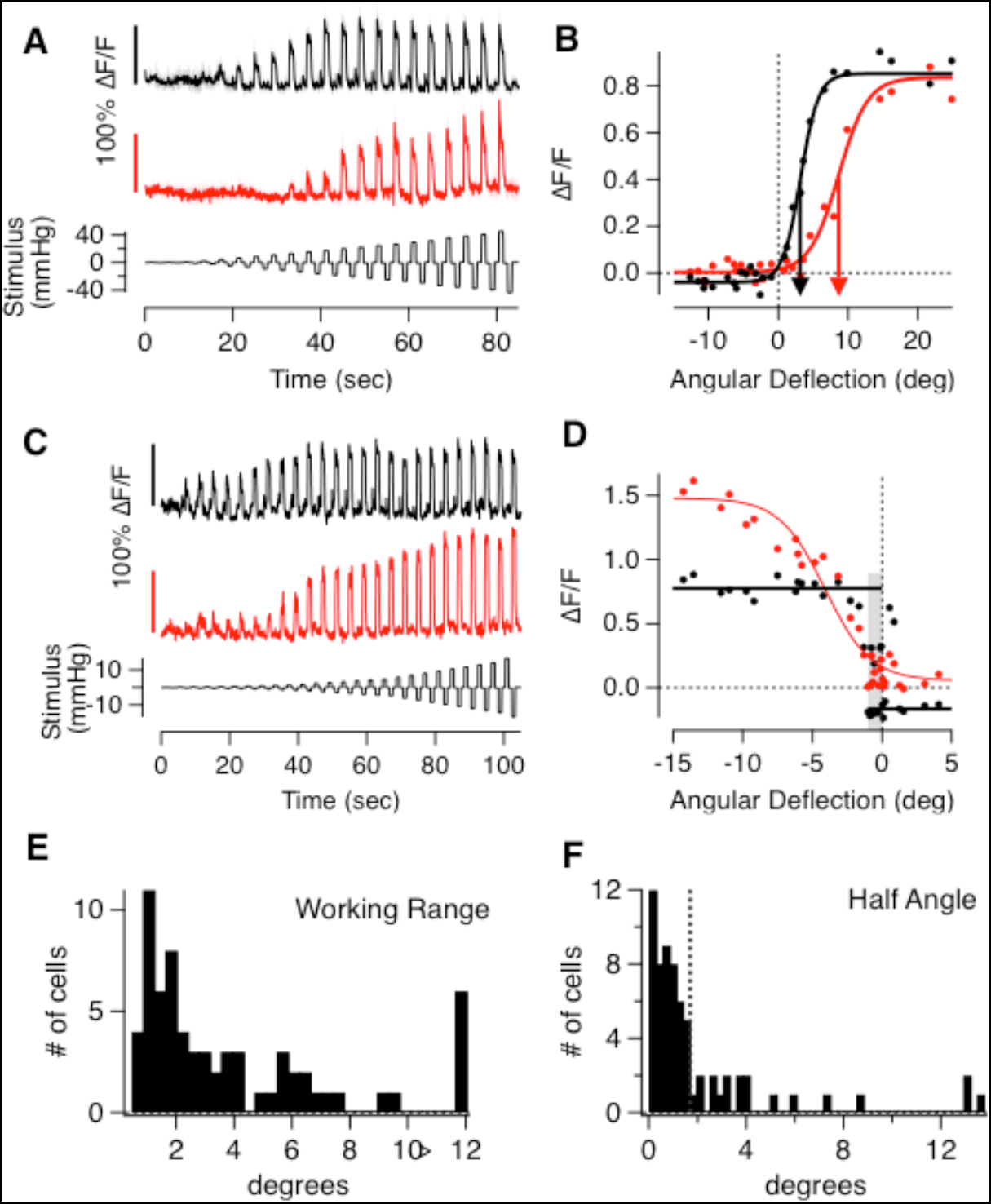
The transfer characteristics of hair cells in the lateral line. **(A)** Responses of two hair cells recorded in the same neuromast, both responding to positive pressure steps. The upper hair cell (black) responds to smaller deflections than the bottom (red). **(B)** Peak iGluSnFR signals (R) from A plotted as a function of the angular deflection of the cupula (X). These stimulus-response relations could be described by a two-state Boltzmann function (equation 1), with parameters R_max_, R_min_, X _1/2_ and X_s_: R_max(1)_ = 0.89 ± 0.02, R_min(1)_= -0.04 ± 0.01, X_1/2(1)_= 3.19 ± 0.15° and X_s(1)_= 1.28 ± 0.15, R_max(2)_ = 0.83 ± 0.02, R_min(2)_= 0.00 ± 0.01, X_1/2(2)_ = 8.65 ± 0.34° and X_s(2)_= 2.07 ± 0.3. **(C)** Responses of two hair cells from another neuromast, which differ more significantly in their working range. **(D)** Stimulus-response relations of the cells in C. The cell depicted in red had a working range of 7°, and the one in black < 1° (grey bar). **(E)** A histogram of working ranges measured in 67 hair cells. About 30% of the cells have working ranges within 1.5°. The last bin contains all cells above 12°. **(F)** Histogram of the X_1/2_ values in 67 hair cells. The majority (75%) had X_1/2_ < 2° (dashed line).

Where R_max_ is the saturating response in the preferred direction, R_min_ is the maximum change in the null direction, X_1/2_ is the rotation that half-activates and X_S_ is the slope factor. Notably, this equation also provides a good description of the relation between hair bundle displacement and mechanotransducer current in hair cells of a number of species, where it has been interpreted as reflecting the transition of the mechanotransducer channel between an open and a closed state (Holton & Hudspeth 1986; Crawford et al 1989).

The two synapses in Fig. 2B differed most significantly in X_1/2_ (arrowed), which is the point at which the gradient of the input-output relation is at its steepest and the sensitivity of the receptor at its maximum (Dayan and Abbott, 2001)Butts & Goldman 2006). The distribution of half-angles across 67 hair cells is shown by the histogram in Fig. 2F. Seventy percent of synapses had half-angles below 2°, corresponding to deflections less than 170 nm at the top of the hair bundle ∼5 μm tall (McHenry et al 2008; Maeda et al 2017). The majority of hair cells in the lateral line therefore have a mechanical sensitivity comparable to auditory hair cells in mice and other species (reviewed by Fettiplace & Kim 2014).

### Heterogeneous transfer characteristics of hair cells within individual neuromasts

The input-output relation of hair cells within a neuromast also varied significantly in their working range (WR) - the deflection required to increase the response from 10% to 90% of maximum (Markin & Hudspeth 1995). The hair cells featured in Fig. 2A and B operated over relatively broad working ranges of 490 nm (5.6°) and 790 nm (9.1°), respectively. Other receptors, however, operated with much narrower working ranges of 90 nm (∼1°), as shown by the black trace in Fig. 2C and the plot in Fig. 2D. Another cell in the same neuromast signalled deflections with a working range of 630 nm (red traces in Fig. 2C and D), again demonstrating the co-existence of receptors with significantly different transfer characteristics. The distribution of working ranges across 67 synapses is shown by the histogram in Fig. 2E, which displayed an initial peak followed by a long tail. The initial peak contained about 60% of synapses and was centred at ∼1.5°, which is equivalent to deflections of 130 nm. These estimates fall within the range of measurements (20-400 nm) made in auditory and vestibular hair cells of a number of species (Fettiplace & Kim 2014). The other 40% of synapses, however, signalled much larger deflections, with working ranges between 5° and 30° (0.5 - 2.5 μm).

The variety of input-output relations measured using iGluSnFR was not expected and we wondered if it might reflect an unexpected property of the reporter. We therefore also assessed the signal transmitted from hair cells by imaging the calcium in the post-synaptic afferent using GCaMP6f under the pan-neuronal *HuC* (*elavl3*) promoter (Experimental Procedures). Fig. 3A-C shows responses in two ROIs within one neuromast and Fig. 3D-F shows the responses in four ROIs of a second. Equation 1 also provided a good description of the input-output relation measured as a post-synaptic calcium signal and the half-angles and working ranges again varied significantly between different synaptic contacts in the same neuromast (Fig. 3G and H). Together, the results in Figs. 2 and 3 demonstrate that the neuromast encodes deflections of the cupula through a mixed population of receptors operating over different ranges.

**Figure 3.**
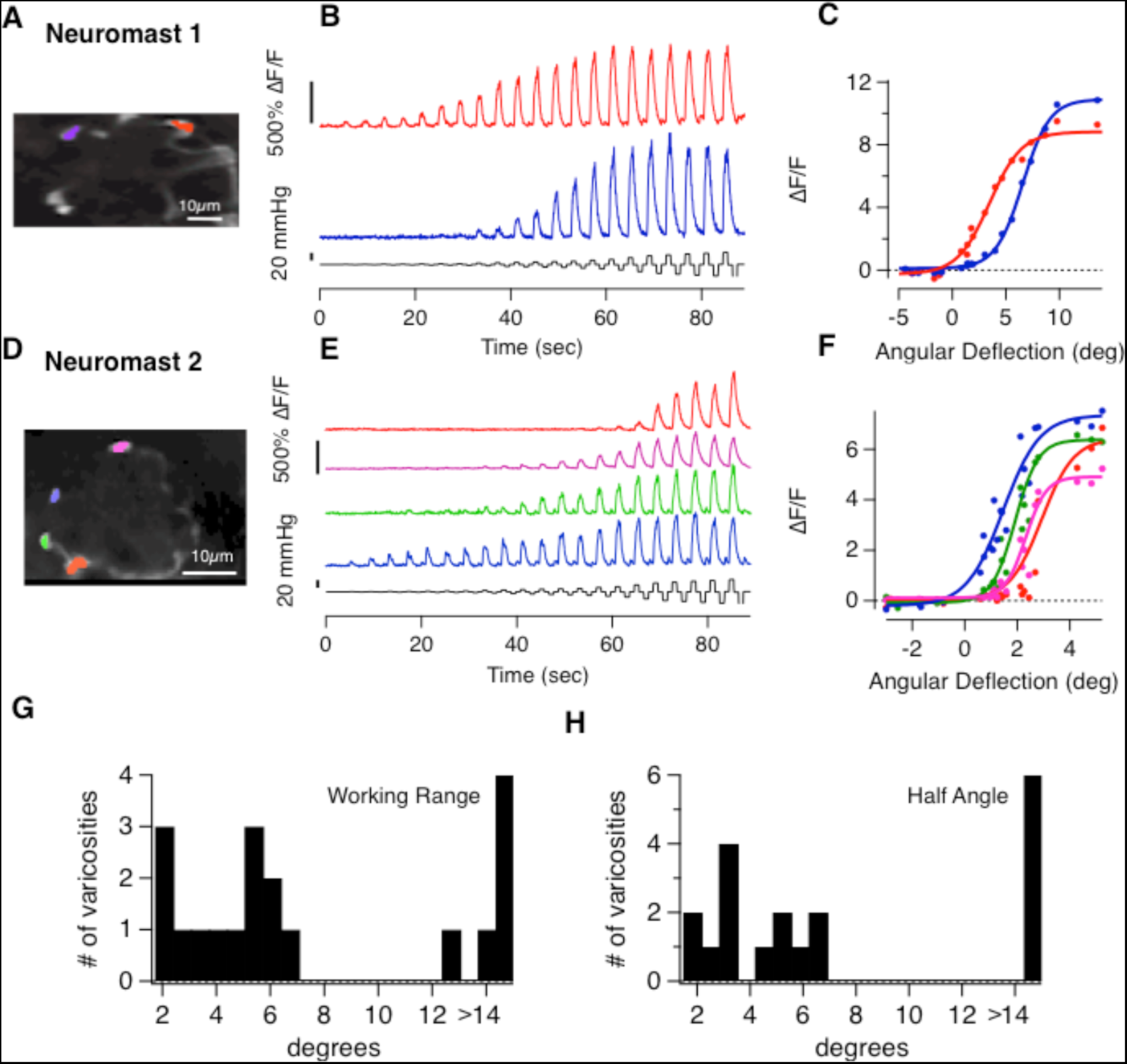
Heterogeneous transfer characteristics of hair cells revealed by measuring calcium signals in afferent neurons. (**A**) The Tg[HuC::GCaMP6f] line expresses the calcium indicator GCaMP6f in afferent neurons of the posterior lateral line but not in hair cells. (**A-C** and**D-F**) show the variety of stimulus response relations from hair cells in two neuromasts. In Neuromast 1 the red and blue traces in (B and C) have an X_1/2_ of 3.5° and 6.6° and a working range of 6.6° and 5.5°, respectively. In neuromast 2 (D-F) X_1/2_ ranges from 1.4° to 3° and the working range spans from 1.7° to 3°. (**G** and**H**) The distribution of working ranges and half-angles in 19 postsynaptic varicosities.

### Individual hair cells can encode opposing directions of motion

The two populations of hair cells polarized to opposite directions allow the neuromast to encode the direction of a stimulus using a “push-pull” system, similar in principle to the ON and OFF channels in the retina (Ghysen & Dambly-Chaudière 2007; Masland 2012). It is less clear whether individual hair cells also encode opposing directions of motion. Afferents leaving the neuromast spike spontaneously and this activity is driven by the release of glutamate from hair cells at rest, which is in turn dependent on the activity of the mechanotransducer channel (Trapani & Nicolson 2011). Can this activity be reduced by deflection in the non-preferred direction? The red trace in Fig. 4A shows an example where this is indeed the case: deflections of the cupula in the non-preferred direction caused a decrease in glutamate release and the input-output relation in Fig. 4B shows that this synapse used 40% of its dynamic range to signal motion in the null direction. In contrast, another hair cell in the same neuromast was completely rectifying, only modulating release in the preferred direction (black trace in Fig. 4A).

**Figure 4.**
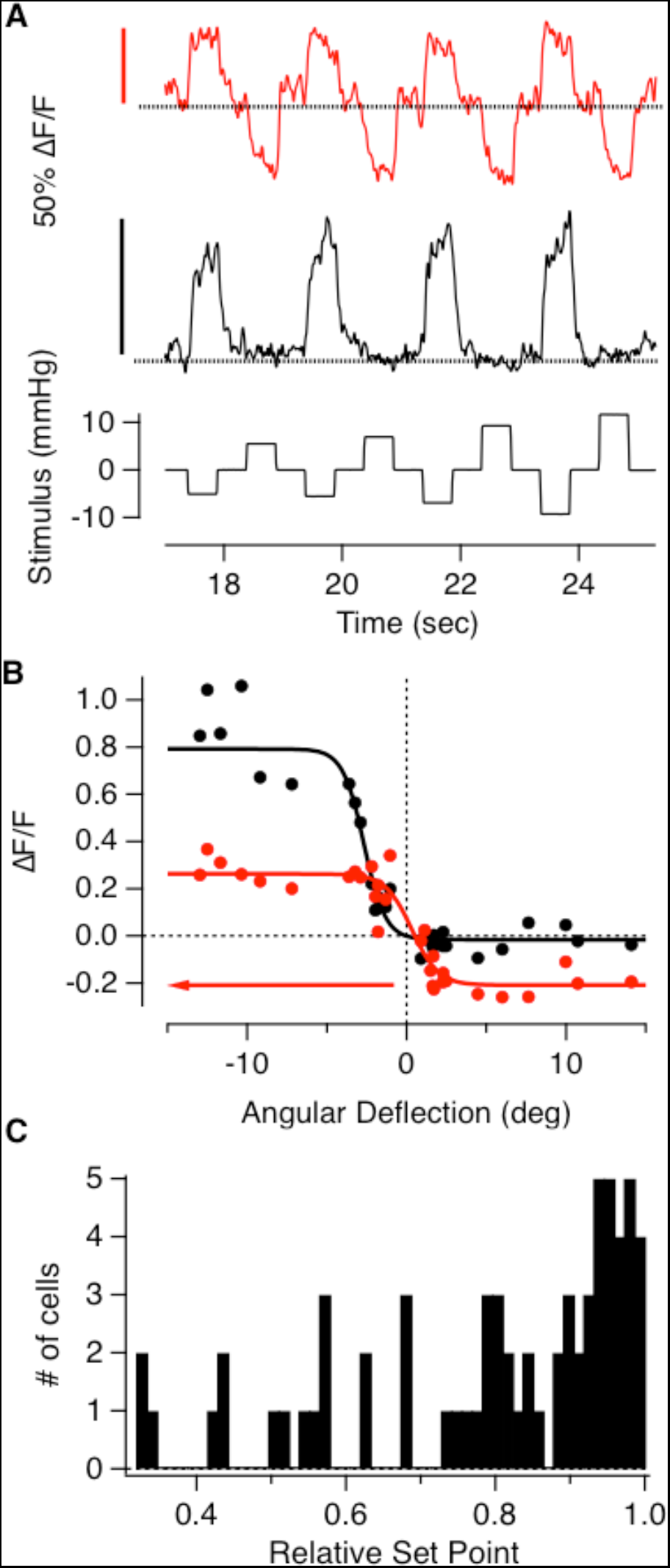
Push-pull signalling in individual hair cells. (**A**) Glutamate release from two hair cells in the same neuromast, both polarized in the negative direction. The cell depicted in black was completely rectifying but the cell in red could also clearly signal deflections in the positive direction as a decrease in glutamate release. (**B**) Stimulus-response relations of the hair cells in A. R_max(1)_= 0.81 ± 0.04, R_min(1)_= -0.02 ± 0.07, X_1/2(1)_= -2.68 ± 0.18° and X_s(1)_= 0.67 ± 0.16, R_max(2)_= 0.47 ± 0.03, R_min(2)_= -0.21 ± 0.05, X_1/2(2)_= 0.32 ± 0.36° and X_s(2)_= 0.83 ± 0.22. (Axis is revevrsed to represent the sensitivity to negative deflections. The relative set-points of the black and red relations were 1 and 0.4, respectively. (**C**) Distribution of the relative set-points from 67 hair cells. Although the majority were strongly rectifying with relative set points close to 1, there was a large degree of variability.

The ability of individual hair cells to signal opposite directions of motion was quantified as the “relative set-point” for glutamate release - the fraction of the total dynamic range of the synapse modulated by deflections from rest in the preferred direction. The distribution of relative set-points across a sample of 67 hair cells is shown by the histogram in Fig. 4C. In 27% of synapses this fraction was less than 0.7, i.e more than 30% of the dynamic range was used to signal deflections in the null direction as a decrease in the rate of glutamate release. These results demonstrate that while the output from most hair cells in the posterior lateral line rectify strongly, the large majority can encode deflections of the cupula both towards the head and away. This property will allow for larger differential signals in the two afferents, whereby an increase in the spike rate of one occurs simultaneously with a decrease in the rate of the second to *below* the spontaneous rate in the absence of a stimulus.

### A mixed population of high- and low-sensitivity hair cells

How do hair cells with different transfer characteristics act as a population to encode a deflection of the cupula? To obtain an overall picture of how the neuromast operates we estimated the total input to a single afferent by averaging the stimulus-response relation from 67 hair cells, assuming that hair cells of opposite polarity were, on average, mirror-images of each other (Fig. 5A). This assumption was based on the observation that the stimulus-response relations of hair cells signalling deflections towards the head were not significantly different from those signalling deflections towards the tail. The tuning curve averaged over all hair cells could be described as the sum of two sigmoid functions with significantly different slope factors and half-angles (X_S(1)_= 0.4 ± 0.1, X_S(2)_ = 1.9 ± 0.9, X_1/2(1)_ =0.6 ± 0.09° and X_1/2(2)_ = 6.1 ± 1.1°). The distribution of half-angles shown in Fig. 3C also indicated two basic populations of hair cells, separable either side of X1/2 = 2°, so we also averaged the stimulus-response relations for hair cells above and below this threshold, as shown in Fig. 5B. The slope factors and half-angles describing the transfer function of these two populations were X_S(<2°)_ = 0.5 ± 0.04, and X_1/2(<2°)_ = 0.7 ± 0.04 (red trace) and X_S(>2°)_ = 1.8 ± 0.3, and X_1/2(>2°)_ =5.0 ± 0 (blue trace). Separating these populations according to X1/2 revealed another important functional difference: hair cells of low sensitivity were completely rectifying with a relative set point of one while cells of high sensitivity had a relative set point of 0.8.

**Figure 5.**
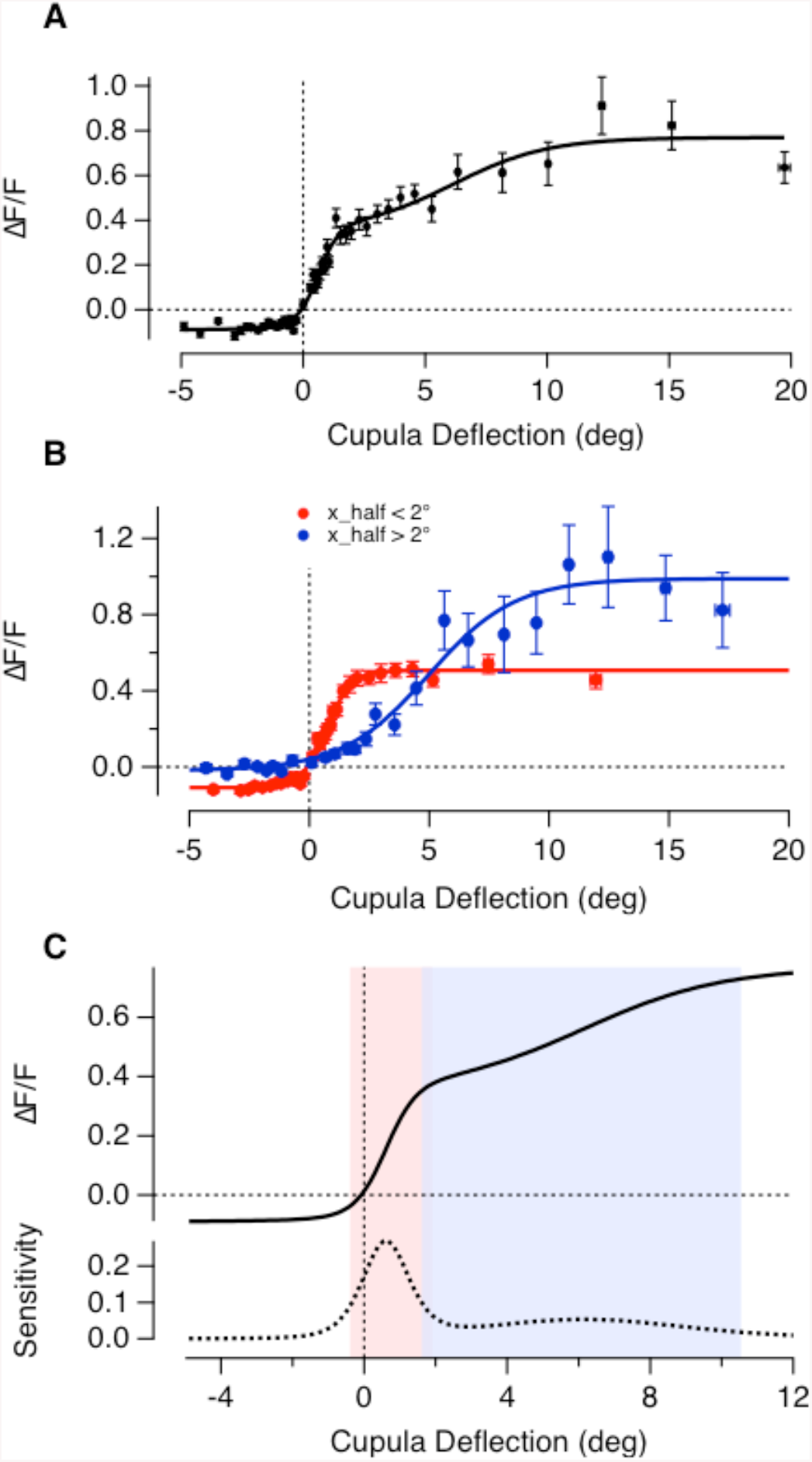
The average transfer characteristics of hair cells in a neuromast. (**A**) 2800 paired measurements of cupula deflection and peak glutamate release, recorded from 67 hair cells. All responses are plotted as a function of the magnitude of deflection, irrespective of direction. A good empirical description was provided by the sum of two Boltzmann relations (equation 1) as shown by the fitted curve (R_min_= 0.1 ± 0.02, R_max(1)_= 0.44 ± 0.09, R_max(2)_= 0.42 ± 0.12, X_1/2(1)_= 0.60 ± 0.1°, X_1/2(2)_= 6.14 ± 1.17°, X_s(1)_= 0.42 ± 0.12, X_s(2)_= 1.95 ± 0.95). The working range of the whole population is 8.9°. The grey curve is the signal predicted to be transmitted to the afferent in the non-preferred direction, assuming that the two groups of hair cell operate symmetrically. (**B**) The average stimulus-response relation of hair cells separated into two groups based on half-angle with a threshold of 2°. The two subsets had average half-angles of 0.7° (red) and 5° (blue). The high- and low-sensitvity groups of hair cell also differed significantly in R_min_, the maximum change in the null direction, with values of -0.11 ± 0.01 and -0.02 ± 0.02, respectively. The working rages of these populations were 2.3° and 8° respectively and overlapped between 1° and 1.9°. (**C**) The sensitivity of the whole population calculated as the derivative of the fit in (A). Small deflections (below 2°) are encoded with high sensitivity by the large population of hair cells whereas the second smaller population extends the dynamic range significantly to also capture larger cupula deflections, ranging beyond 10°.

The sensitivity of a sensory system can be quantified as the change in response per unit change in stimulus i.e the first derivative of the stimulus-response relation (Dayan & Abbott 2001). The dashed line in Fig. 5C shows this function for the average output of all 67 hair cells, from which three features stand out. First, the neuromast achieves maximum sensitivity at deflections of just ∼40 nm at the tip of the hair bundle. Second, deflections from rest can be signalled with a sensitivity about 63% of maximum, either as an increase or a decrease in glutamate release, and this is made possible by the relative set-point of the high-sensitivity population of hair cells. Third, the high-sensitivity population saturates at deflection of ∼220 nm, but the dynamic range of the neuromast as a whole is extended up to about 1 μm by the low-sensitivity population. Acting together, these two groups of hair cells will make the neuromast very sensitive to small deflections of the cupula while maintaining a large dynamic range.

### Heterogeneous adaptive properties of hair cells within individual neuromasts

A second strategy by which sensory systems prevent saturation is adaptation, a time-dependent change in gain that adjusts the working range in response to the recent history of stimulation (Wark et al 2007). In the lateral line, ramped deflections of the cupula cause the spike rates in the afferents to adapt strongly (Haehnel-Taguchi et al 2014) but the cellular mechanisms are not clear. In hair cells of the auditory and vestibular systems the two main loci of adaptation are the mechanoelectrical transducer (MET) in the hair bundle (Howard & Hudspeth 1987; Shepherd & Corey 1994;(Holt et al., 2002);(Kennedy et al., 2003)), reviewed by (Ricci & Kachar 2007) and depression at the ribbon synapse (Schnee et al 2005; Schnee et al 2011; Goutman 2017). These two processes have been studied in isolation, but little is known about how they act together to adjust the input-output relation of the hair cell.

To investigate the adaptive properties of the neuromast we applied saturating or near-saturating pressure steps of 2 s or more. In 61 of 65 hair cells adaptation was apparent as synaptic depression but there was a large degree of variability within the same neuromast. For instance, Fig. 6A shows an example in which glutamate release fell to about 30% of peak in one hair cell (red) but only to 70% of peak in another (black). We quantified the reduction in glutamate release as an adaptation index (AI), which varied from 1 (complete recovery of the iGluSnFR signal) through zero (no change) to negative values (reflecting an acceleration of glutamate release; Experimental Procedures). The distribution of AIs measured at the end of a saturating 5 s pressure step varied widely, as shown by the distribution in Fig. 6C. The speed of decay of the response could be described by a time-constant of 2 s or less in 74% of cells, and the distribution in this subpopulation is shown in Fig. 6D. The remaining cells had a decay time-constant greater than 2 s that could not be estimated reliably from a 5 s record. The smallest time constant we observed was 130 ms and 30% of all cells had a decay constant below 500 ms.

**Figure 6.**
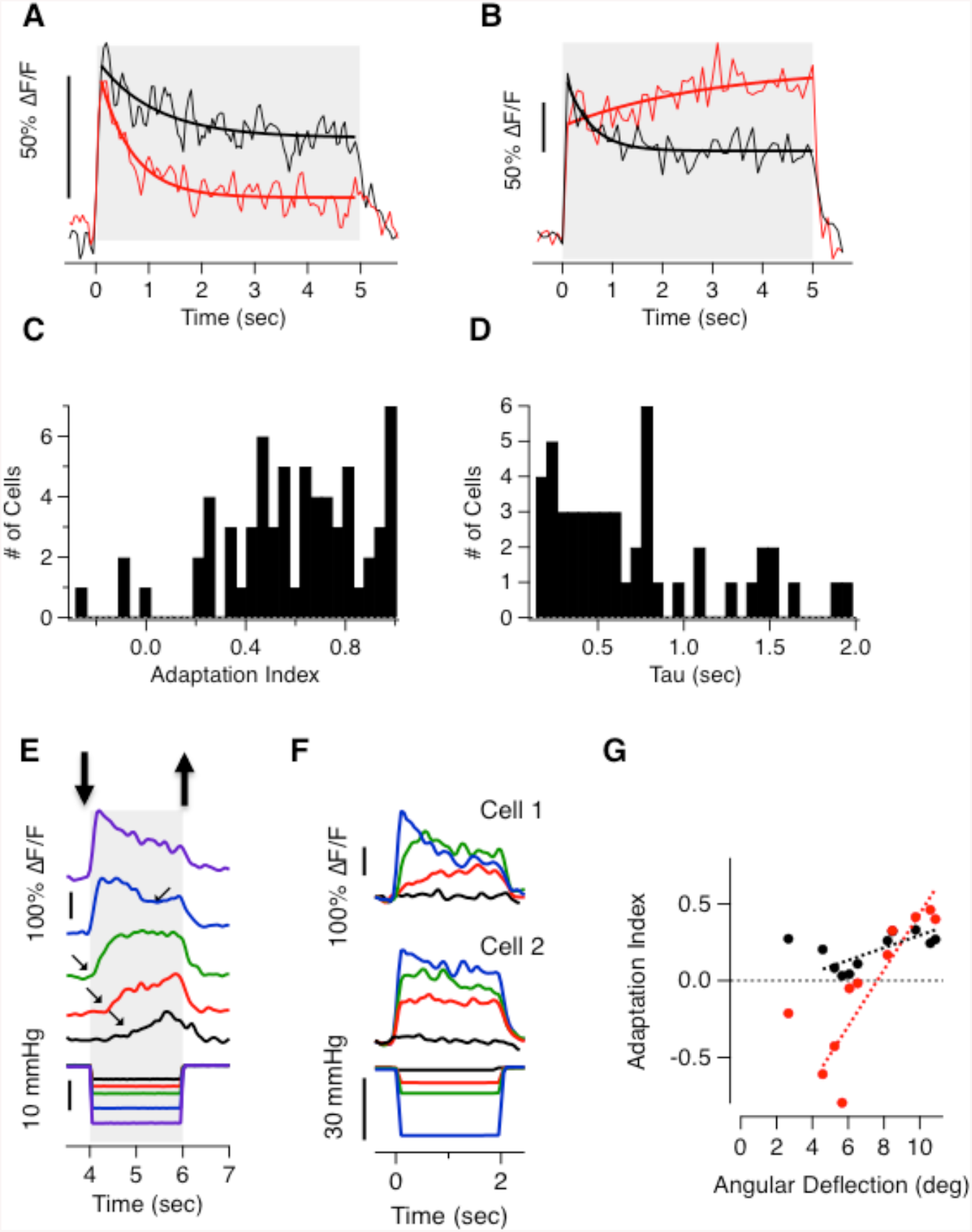
Heterogeneous adaptive properties of hair cells. **(A)** Two hair cells from the same neuromast adapt to different extents to a 5 s step deflection. The red cell adapted with a time-constant of 0.61 ± 0.09 s (solid line) with an adaptive index of 0.65. The black cell adapted with a time-constant of 1.09 ± 0.32 s with an adaptive index of 0.42. **(B)** The iGluSnFR signal from two hair cells of a second neuromast. Note that while one cell adapted (black, AI = 0.53) the second sensitized (AI = -0.13). **(C)** Distribution of adaptation index from 65 hair cells stimulated with a 5 s step. **(D)** Distribution of the decay time constants from the 55% of cells that could be fit with tau < 2 s. In the remainder of cells tau was greater than 2 s and could not be estimated. (**E**) A series of iGlusnFr signals from one hair cell. Large deflections cause glutamate release to rapidly reach a peak and then depress while small deflections do not generate a detectable signal for ∼0.5 s after which glutamate release accelerates. (**F**) Two hair cells from the same neuromast responding to negative deflections. Cell 1 shows sensitization to small deflections but cell 2 does not. (**G**) The adaptation index as a function of deflection angle for cell 1 (red) and cell 2 (black) from B. Dashed lines are fits to the points.

These results demonstrate that hair cells of the lateral line apply different temporal filters: in some, a step stimulus generates a transient response that signals most strongly the onset of the deflection (e.g red trace in Fig. 6A) while in others the output is more sustained and effectively signals the duration (e.g black trace in Fig. 6B). An analogy can again be made with the transformation of signals in the retina, where the distinction between transient and sustained neurons has long been recognized (Baccus 2007; Masland 2012).

### Hair cells sensitized by small deflections

Adaptation in sensory systems is usually thought of as a reduction in gain acting to prevent saturation (Wark et al 2007), but in 3 cells a strong deflection caused the rate of glutamate release to *increase* over periods of a few seconds, as shown by the red trace in Fig. 6B. The ability of some hair cells to sensitize to a constant stimulus became more apparent when measuring responses to smaller deflections. For example, Fig. 6E shows the iGluSnFR signal from a single hair cell in which relatively small deflections did not generate a significant output within the first 500 ms, after which glutamate release gradually accelerated (black and red traces). A deflection of intermediate amplitude generated a response that gradually achieved a steady state (green) while larger deflections generated responses that reached a peak within ∼200 ms and then depressed strongly (blue and violet traces). The ability to sensitize to small deflections was also one that varied between different hair cells within one neuromast, as shown by the two examples in Fig. 6F. A shift from sensitization to depression with increasing stimulus strength was observed in 4 of 14 hair cells stimulated for 2 s and could be described by the AI plotted as a function of deflection angle, as shown in Fig. 6G.

A functional role for sensitization can be appreciated by asking when the offset of the stimuli in Fig. 6E are signaled most clearly: the largest decrease in glutamate release occurred after *smaller* deflections that caused sensitization (red and green traces). A rapid sensitization of subsets of neurons has also been observed in the retinal response to an increase in contrast (Kastner & Baccus 2011; Nikolaev et al 2013; Kastner & Baccus 2013), where the mixture of depressing and sensitizing units allows the retinal circuit to signal natural fluctuations in contrast more effectively than would be the case if all neurons simply reduced their gain when contrast increased (Kastner & Baccus 2011). A qualitatively similar mechanism appears to operate in the neuromast where depressing synapses most clearly signal a deflection away from rest and sensitizing synapses signal a deflection back to rest.

### Population signalling of a return to rest

Here we have shown that push-pull signalling within a neuromast is facilitated by the high-sensitivity population of hair cells with relative set-points below 1 (Fig. 4A) and by a mixture of sensitizing and desensitizing hair cells (Fig. 6). A third and distinctive property of the population signal is shown by the pair of synapses in Fig. 7A, which were of opposite polarity and completely rectifying such that a deflection in the non-preferred direction did not generate a response (green boxes). A return to rest from the non-preferred direction did, however, generate a strong and transient release of glutamate (black boxes). In other words, only one of the two hair cells signalled a deflection from rest (green boxes), while *both* signalled a recovery to rest with transient responses in opposite directions (black boxes). Such a “reset” or “rebound” signal was observed in 33 of 55 hair cells stimulated with steps of 1 s duration.

**Figure 7.**
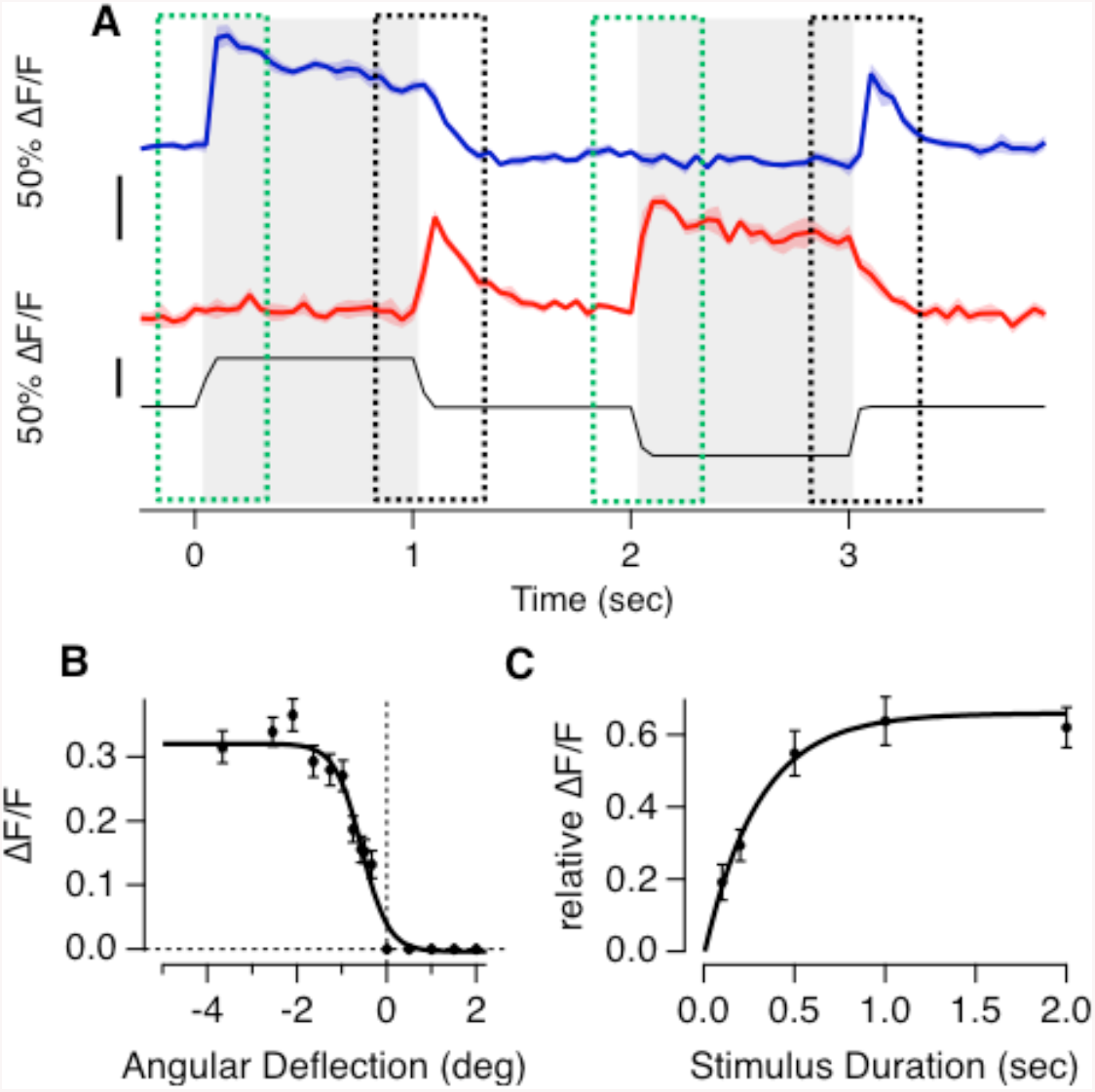
Hair cells signalling a return to rest. **(A)** Responses of two hair cells that generate a ‘reb^o^ und-response’ – a large and transient release of glutamate after the release of a deflection in the null direction. **(B)** The relationship between angular deflection in the null direction and amplitude of the rebound response. Data is averaged from 33 hair cells in which the largest rebound response exceeded 20% of the maximum response in the preferred direction (R_max_= 0.32 ±0.01, R_min_= 0 ± 0.01, X_1/2_= -0.56 ± 0.05 and X_s_= 0.3 ± 0.05). **(C)** The relationship between duration of deflection in the null direction and the magnitude of the rebound response (as a fraction of the response to stimulation in the preferred direction). These experiments were performed with large deflections generating the maximum rebound. Results are described by an exponential that yields a time constant of 0.3 sec for the development of the rebound response.

The amplitude of the reset signal depended on both the magnitude and duration of the preceding deflection in the non-preferred direction. The dependence on magnitude could be described by a two-state Boltzmann equation with a half-angle of -0.6° (50 nm; Fig. 7B) and during a large deflection the response developed with a time-constant of 0.3 s (Fig. 7C). The reset signals transmitted through the hair cell synapse are qualitatively similar to the process of “negative adaptation” described in the MET current of a variety of hair cells and is therefore likely driven by these channels (Holt et al 2002; Hirono et al 2004; Stauffer et al 2005; Hudspeth, 2014). Negative adaptation within the neuromast generates a population signal that encodes the cessation of a stimulus, as would occur, for instance, in the intervals between swimming bouts (Russell & Roberts 1972; Palmer et al 2005). The growth of the reset signal with time and angle of deflection indicates that it might also be used to integrate flow gradients against the side of the fish, the detection of which underlies rheotaxis (Oteiza et al 2017).

## Discussion

Sensory processing begins with the detection and encoding of different features of the input arriving at the peripheral sense organ. This study demonstrates that neuromasts of the lateral line encode a mechanical stimulus through a number of filters that vary in sensitivity, polarity and temporal and adaptive properties. Qualitatively similar filters operate in, for instance, the retina (Baccus 2007; Masland 2012; Kastner & Baccus 2011) but, while in the neuromast they all reside in the primary receptors, in the retina they are also functions of the downstream circuitry. Directly measuring the mechanical stimulus, deflection of the cupula, and the synaptic output, release of glutamate, has allowed us to specify how the transfer characteristics of the hair cells within a neuromast vary and here we discuss how these variations contribute to the information that can be transmitted as well as the cellular mechanisms by which they might arise.

### Sensitivity, working range and set-point

The high-sensitivity group of hair cells operated with half-angles around 1.5° (Fig. 2F), which corresponds to a displacement of 130 nm at the tip of the stereocilia and is comparable to the sensitivity of auditory and vestibular hair cells in mice and other species (Fettiplace & Kim 2014). This group saturated at deflections of ∼220 nm (Fig. 5B), which is much narrower than the working range of the neuromast as a whole (Haehnel-Taguchi et al 2014). The difference can be accounted for by the second, low-sensitivity population of hair cells that extended the dynamic range of the neuromast beyond 1 μm. A strategy for detecting stimuli with high- and low-sensitivity receptors allows the neuromast to encode weak stimuli effectively without saturating.

An important feature of the high-sensitivity hair cells was a relative set-point that allowed small deflections in either direction to modulate glutamate release (Figs. 4 and 5). The sensitivity to deflections from rest achieved 63% of the maximum measured at ∼40 nm (Fig. 5C) so it seems likely that deflections less than 50 nm will be transmitted to targets in the hindbrain. The behavioural significance of detecting such small deflections is, however, harder to judge and would require one to measure (or perhaps calculate) deflections of the cupula in a motile fish. One possibility is that the high-sensitivity group of hair cells are involved in the sensing of flow velocity gradients around the body of the fish that have recently been shown to underly rheotaxis in the absence of visual input (Oteiza et al 2017). Low-sensitivity hair cells would then be available to encode stronger stimuli triggering reflexes such as the escape response.

What are the mechanisms that underly these variations in sensitivity? One potential locus is the transduction process within the hair bundle. The relationship between displacement from rest and the MET current has also been found to vary in different hair cells recorded within a single experiment in the auditory and vestibular systems (Holt et al 1997; Stauffer & Holt 2007). The working range assayed using the MET current can be related to the force required to open the MET channel and, therefore, to the stiffness of the “gating spring” that pulls on the channel when the hair bundle is deflected in the preferred direction (Howard & Hudspeth 1987). Models based on variations in the stiffness of the gating spring can account for adaptive changes in the working range and set-point and are consistent with the correlation that we observed between higher sensitivity and positive set-point (Fig. 5B). A stiffer gating spring would cause some of the MET channels to be open at rest as well as a larger increase in open probability for a given deflection. Changes in stiffness have been suggested to reflect actions of calcium at the MET channel or motions of the tip-links mediated by myosin1c (Hacohen et al 1989; Holt et al 2002; Hirono et al 2004; Stauffer et al 2005), but the molecular mechanisms are still debated (Peng et al 2013; Fettiplace & Kim 2014).

A change in sensitivity measured at the output might also reflect the process of Ca^2+^-triggered exocytosis at the active zone. Push-pull modulation of glutamate within individual hair cells requires that the resting potential sits in a range where CaV1.3 channels under the ribbon are activated sufficiently to drive vesicle fusion (Platzer et al 2000; Olt et al 2014). If the resting potential is hyperpolarized relative to the threshold for Ca^2+^ channel activation the output of the hair cell will be completely rectifying, as observed in the low-sensitivity population of hair cells. Conductances that might cause resting potentials to vary include K^+^ channels in the basolateral membrane (Olt et al 2014) and Ih inward rectifiers (Trapani & Nicolson 2011). The voltage-dependence of Ca_V_1.3 channels may also be subject to modulation that alters the threshold for activation and the steepness of the current-voltage relation (Striessnig et al 2010). Even within a single hair cell active zones may vary in the number of calcium channels and their voltage-dependence, leading to different release rates driven by the same receptor potential (Ohn et al 2016). Finally, variations in sensitivity at the output might also reflect differences in the efficiency with which a given Ca^2+^ signal triggers exocytosis (Olt et al 2014).

### Adaptation

We also found a large variability in the adaptive properties of hair cells within individual neuromasts, in terms of the degree, speed and even direction (Fig. 6). Such a mixture of responses will help the population within a neuromast to signal both sustained stimuli, such as water flow (Voigt et al 2000), and sudden deviations, such as eddy currents or the motion of other creatures in their immediate environment (McHenry et al 2009).

Adaptation of the MET current varies strongly between hair cells but can generally be described by two time constants, τ_fast_ in the ms range and τ_slow_ in the tens of ms range (Shepherd & Corey 1994; Holt et al 1997; Vollrath & Eatock 2003; Stauffer & Holt 2007). These processes are, however, orders of magnitude faster than the kinetics of adaptation that we were able to observe at the output of the synapse (Fig. 6), indicating that the rate-limiting process is downstream of the MET channel. A strong possibility is depletion of releasable vesicles at the ribbon synapse. Indeed, capacitance measurements of exocytosis in hair cells of the lateral line demonstrate that a strong step depolarization stimulates release that decays with time constants of ∼500 ms (Lv et al 2016; Eatock 2000), while optical measurements at ribbon synapses in the retina demonstrate that these can also depress with time-constants of a few seconds (Nikolaev et al 2013).

An intriguing finding was that a small number of hair cells sensitized during small deflections by gradually increasing the rate of glutamate release over periods of hundreds of milliseconds (Fig. 6). As well as signalling weak but continuous stimuli, sensitization will cause the offset to be signalled more strongly than the onset. Both these phases of the output may play a role in behaviours such as rheotaxis, where constant but slow flows of water deflect the neuromasts for prolonged period, and deviations from this flow orientation trigger a righting behavior (Oteiza et al 2017). Studies of the MET channel do not usually test its modulation over these longer time-scales, so it is difficult to assess if these might be the cause of sensitization. A stronger candidate for the intracellular signal integrating these weak stimuli is the accumulation of calcium in the cytoplasm, which has been shown to accelerate the trafficking of vesicles for release (Castellano-Muñoz & Ricci 2014) as well as exocytosis itself (Pangršič et al 2015). This replenishment process will likely also determine the degree of adaptation achieved in the steady state (Baden et al 2014).

A second process that signalled the offset of a stimulus was so-called “negative adaptation” to hair bundle deflections away from the kinocilium, which primed the hair cell to generate a transient burst of glutamate release when the cupula returned towards rest (Fig. 7). The amplitude of this signal was a function of both the amplitude and duration of the deflection and therefore encoded the integrated stimulus. A signal that specifically informs upstream circuits of the cessation of an input might be used to, for instance, readjust the gain with which incoming signals control motor behaviour.

## Experimental Procedures

### Fish husbandry

Adult zebrafish (Danio rerio) were maintained in fish water at 28.5°C under a 14:10 hour light:dark cycle under standard conditions (Brand et al 2002). Fish were bred naturally and fertilized eggs were collected, washed with distilled water and transferred into 50 ml of E2 medium (concentrations in mM: 0.05 Na2HPO4, 1 MgSO4 7H2O, 0.15 KH2PO4, 0.5 KCl, 15 NaCl, 1 CaCl2, 0.7 NaHCO3, pH7-7.5). At 24 hours post fertilisation (hpf) 1-phenyl2-thiourea (pTU) was added to yield a final concentration of 0.2 mM to inhibit pigment formation. All procedures were in accordance with the UK Animal Act 1986 and were approved by the Home Office and the University of Sussex Ethical Review Committee.

### Fish Lines

The Tg[HuC::GCaMP6f] line (kindly provided by Isaac Bianco) expresses GCaMP6f pan-neuronally, including in afferent neurons of the lateral line, but not in hair cells. The Tg[Sill2, UAS::iGluSnFR] line expresses iGluSnFR (Marvin et al 2013) exclusively in afferent neurons of the lateral line system (Pujol-Martí et al 2012). It was generated by injecting the Sill2 construct, containing the Sill enhancer driving expression of the Gal4-VP16 element (kindly provided by Hernan Lopez-Schier), into single-cell stage embryos expressing 10xUAS::iGluSnFR. Larvae were sorted for expression of iGluSnFR and reared to adulthood to identify fish with germ-line transmission (founders). To generate the Tg[Sill2, UAS::iGluSnFR, Rib::Rib-mCherry], the Sill construct was injected into the embryos derived from an outcross between 10xUAS::iGluSnFR and Rib::Rib-mCherry (Odermatt et al 2012) adults and screened for the expression of iGluSnFR and mCherry. This line allowed us to image the iGluSnFR signal in afferent neurons while visualising synaptic ribbons in hair cells. All fish were maintained in a nacre mutant background (Lister et al 1999).

### Mounting and Cupula Staining

All experiments were performed at room temperature (20°C – 25°C) using larvae at 7-10 days post fertilisation (dpf). At 4 dpf, embryos were screened for the strongest expression of the appropriate. Larvae were anaesthetised by immersion in 0.016% tricaine (MS-222), diluted in E2. They were then placed side-down into a ‘fish-shaped’ pit carved into a thin (∼1mm) layer of PDMS (Sylgard184, Dow Crowning) on a coverslip. Mechanical stability was provided by a ‘harp’ (Warner Instruments) placed on top of the larva. The pressure applied by the nylon strings was adjusted to allow normal blood flow while maintaining enough pressure to hold down the larvae. The larva was then paralyzed by injection of 0.25mM alpha-Bungarotoxin (αBTX; Tocris) into the heart. To avoid damaging the cupula, special care was taken not to touch the upward-facing side of the fish during the mounting procedure. The cupula was then stained by incubating the fish in a 1:500 dilution of 1mg/ml WGA AlexaFluor-594 or WGA AlexaFluor-350 (Life Technologies) for 2 minutes followed by thorough washing with E2. When counter-staining hair cells with FM4-64 (Synaptored, Biotium) larvae were incubated in a 1 μM solution for 1 minute and then washed with E2.

### Mechanical stimulation

Pressure steps were applied to a neuromast through a glass pipette attached to a high speed pressure clamp (HSPC-1, APA scientific) (Trapani et al 2009). The output pressure was controlled and recorded using mafPC software (courtesy of M. A. Xu-Friedman) running in IgorPro (Wavemetrics) and synchronised to image acquisition. The micropipette was pulled to a diameter of ∼30 μm and the tip bent through 30° using a micro forge (Narishige) to allow liquid flow parallel to the body of the larva. The tip was positioned ∼20 μm above the body, ∼100 μm from the neuromast. This study was confined to neuromasts of the posterior lateral line with an ‘anterior-posterior’ axis of sensitivity. The direction of the pipette (pointing towards the tail or towards the head) was changed during the course of some experiments but did not affect measurements.

### *In Vivo* Two-Photon Imaging

Fish were imaged on a custom built two-photon microscope driven by a mode-locked Titanium-sapphire laser (Chameleon 2, Coherent) tuned to 915 nm. Excitation was delivered through a 40x water immersion objective (Olympus, 40x LUMIPlanF, NA: 0.8) and emitted photons were collected both through this objective and an oil condenser (NA 1.4, Olympus) below the sample. Green and red fluorescence was separated by a 760dcxru dichroic and filtered through 525/70 nm and 620/60 nm emission filters before being focused onto GaAsP photodetectors (Hamamatsu). The microscope was controlled by ScanImage v3.8 (Vidrio Technologies) and image acquisition was synchronised with the stimulus. Movies were acquired at 10-50 Hz.

### Measuring deflections of the cupula

The angular deflection of the stained cupula was assessed by measuring its translational displacement in multiphoton images in planes at different z distances from the surface of the hair cells. These measurements were made at a variety of stimulus pressures and repeated in 3-4 planes 5 μm increments. The central position of the cupula within each frame was extracted by first thresholding the image and then fitting an ellipse to estimate the centre of mass. Next, the translational deflections induced by the applied pressure steps in each plane were calculated: these were consistent with the proximal regions of the cupula (z < 15-20 μm) behaving as a pivoting beam (McHenry & van Netten 2007). In this way, a calibration of the angular deflection of the cupula for each stimulus pressure was obtained for each experiment (Fig. S1). This calibration was repeated if, for instance, the pipette was moved.

### Image Analysis

Images were analysed in Igor Pro (Wavemetrics) using custom-written software including the SARFIA toolbox (Dorostkar et al 2010). Movies containing small drifts in the x-y dimension were registered but movies mages with large drifts, including potential z-motions, were discarded. Regions of Interest (ROIs) were determined using an algorithm that began by identifying pixels with both large signals and high degrees of temporal correlation. Pixels surrounding thee ROI “seeds were added to the ROI until the correlation value fell below a threshold. Background fluorescence was then subtracted and baseline fluorescence (F_o_) was defined as the average fluorescence preceding the first stimulation interval. The change in fluorescence relative to baseline (ΔF/Fo) was calculated and used for further analysis.

The adaptation index (AI) was calculated as:

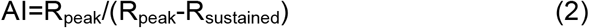

Where R_peak_ is the instantaneous peak response after stimulus onset R_sustained_ is the average responses during the last 100 ms of stimulus presentation. Therefore, an AI close to 0 indicates no adaptation, an AI close to 1 indicates complete adaptation and a negative AI indicates sensitization.

The relative set point (SPr) was calculated as:

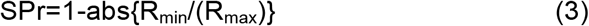

Where R_max_ is the saturating response in the preferred direction of deflection and Rmin is the maximum change in the opposite direction.

## Supplementary Information

### Figure S1. Quantifying the angular deflection of the cupula

Related to Figure 1. Illustrates how the cupula behaved as a rigid beam and how the rotation of that beam around its base was imaged and calculated.

**Supplementary Figure 1.**
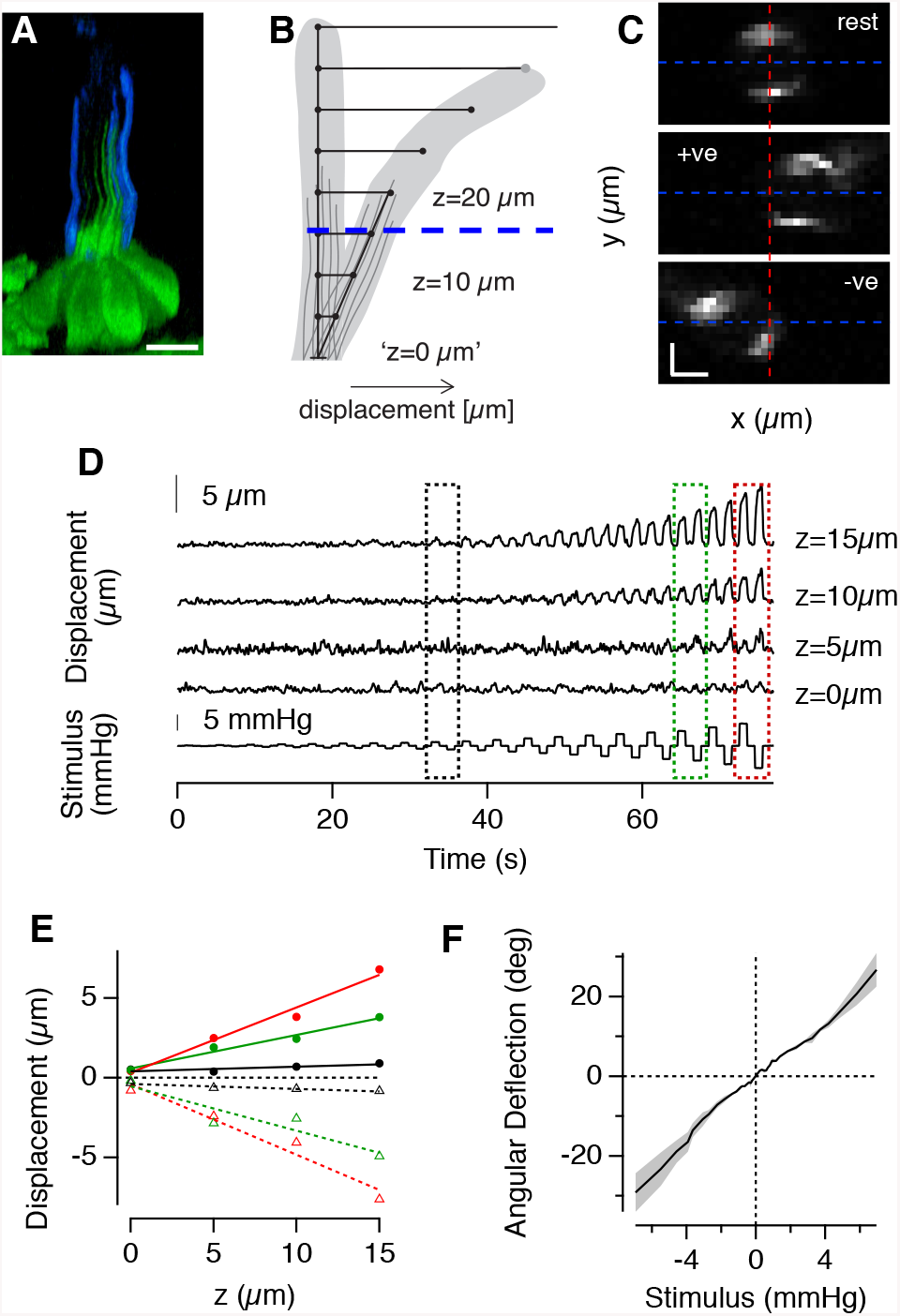
Calibrating the relationship between stimulus pressure and deflection of the cupula. (**A**) Side view of a NM of expressing GFP in hair cells with the surface of the cupula stained with Alexa-350-WGA. Notice how the kinocilia extend roughly half way up the cupula (scale bar 10 μm). (**B**) Schematic of the model used to calculate angular deflections of the cupula from a pivot at its base. The translational deflection for a given stimulus pressure was measured at several distances z above the apical surface of the hair cell. (**C**) Three representative frames of the stained cupula at z = 15 μm at rest and deflected by a positive and negative pressure step.**(D)** Traces showing x displacement as a function of time at four different z distances and for a variety of pressure steps (bottom trace). The movies were obtained at 20 Hz.**(E)** The x displacements to positive and negative pressure steps of increasing magnitude (black, green and red, corresponding to stimuli delivered boxes shown in D). Note that x displacement is directly proportional to z at any pressure, indicating that the cupula moved through a fixed angle, acting as a beam. The angular deflection was calculated as tan^-1^ (Δ/z), where (Δx) was the translation in the centre of mass of the staining in C.**(F)** A calibration curve relating the stimulus pressure to the angular deflection from the measurements in C-D (grey shading is the s.e.m). The relation is roughly linear.

## References

Ayali, A., Gelman, S., Tytell, E.D. & Cohen, A.H. (2009). Lateral-line activity during undulatory body motions suggests a feedback link in closed-loop control of sea lamprey swimming. Can. J. Zool.87(8), 671–83.

Baccus, S.A. (2007). Timing and computation in inner retinal circuitry. Annual Review of Physiology. 69, 271–90.

Baden, T., Nikolaev, A., Esposti, F., Dreosti, E., Odermatt, B. & Lagnado, L. (2014). A synaptic mechanism for temporal filtering of visual signals. PLoS Biol. 12(10), p. e1001972.

Beutner, D., Voets, T., Neher, E. & Moser, T. (2001). Calcium dependence of exocytosis and endocytosis at the cochlear inner hair cell afferent synapse. Neuron. 29(3), 681–90.

Brand, M., Granato, M. & Nüsslein-Volhard, C. (2002). Keeping and raising zebrafish, in Zebrafish: A Practical Approach,Oxford University Press. Oxford, UK, 7–37.

Brandt, A., Khimich, D. & Moser, T. (2005). Few CaV1.3 channels regulate the exocytosis of a synaptic vesicle at the hair cell ribbon synapse. The Journal of Neuroscience. 25(50), 11577–85.

Butler, J.M. & Maruska, K.P. (2016). The Mechanosensory Lateral Line System Mediates Activation of Socially-Relevant Brain Regions during Territorial Interactions. Frontiers in Behavioral Neuroscience. 10, p. 93.

Butts, D.A. & Goldman, M.S. (2006). Tuning curves, neuronal variability, and sensory coding. PLoS Biol. 4(4), p. e92.

Castellano-Muñoz, M. & Ricci, A.J. (2014). Role of intracellular calcium stores in hair-cell ribbon synapse. Front Cell Neurosci. 8, p. 162.

Corns, L.F., Johnson, S.L., Kros, C.J. & Marcotti, W. (2014). Calcium entry into stereocilia drives adaptation of the mechanoelectrical transducer current of mammalian cochlear hair cells. Proceedings of the National Academy of Sciences of the United States of America. 111(41), 14918–23.

Crawford, A.C., Evans, M.G. & Fettiplace, R. (1989). Activation and adaptation of transducer currents in turtle hair cells. The Journal of Physiology. 419, 405–34.

Dayan, P. & Abbott, L.F. (2001). Theoretical neuroscience.Cambridge, MA: MIT Press.

Dorostkar, M.M., Dreosti, E., Odermatt, B. & Lagnado, L. (2010). Computational processing of optical measurements of neuronal and synaptic activity in networks. J Neurosci Methods. 188(1), 141–50.

Eatock, R.A. (2000). Adaptation in hair cells. Annual review of neuroscience. 23, 285–314.

Eatock, R.A., Corey, D.P. & Hudspeth, A.J. (1987). Adaptation of mechanoelectrical transduction in hair cells of the bullfrog’s sacculus. The Journal of Neuroscience. 7(9), 2821–36.

Erickson, T., Morgan, C.P., Olt, J., Hardy, K., Busch-Nentwich, E., Maeda, R., Clemens, R., Krey, J.F., Nechiporuk, A., Barr-Gillespie, P.G., Marcotti, W. & Nicolson, T. (2017). Integration of Tmc1/2 into the mechanotransduction complex in zebrafish hair cells is regulated by Transmembrane O-methyltransferase (Tomt). eLife, 6.

Faucherre, A., Pujol-Martí, J., Kawakami, K. & López-Schier, H. (2009). Afferent neurons of the zebrafish lateral line are strict selectors of hair-cell orientation. PloS One. 4(2), p. e4477.

Fettiplace, R. & Kim, K.X. (2014). The physiology of mechanoelectrical transduction channels in hearing. Physiological Reviews. 94(3), 951–86.

Ghysen, A. & Dambly-Chaudière, C. (2007). The lateral line microcosmos. Genes Dev.21(17), 2118–30.

Goutman, J.D. (2017). Mechanisms of synaptic depression at the hair cell ribbon synapse that support auditory nerve function. Proceedings of the National Academy of Sciences of the United States of America. 114(36), 9719–24.

Graydon, C.W., Manor, U. & Kindt, K.S. (2017). In Vivo Ribbon Mobility and Turnover of Ribeye at Zebrafish Hair Cell Synapses. Scientific Reports. 7(1), p. 7467.

Hacohen, N., Assad, J.A., Smith, W.J. & Corey, D.P. (1989). Regulation of tension on hair-cell transduction channels: displacement and calcium dependence. The Journal of Neuroscience. 9(11), 3988–97.

Haehnel-Taguchi, M., Akanyeti, O. & Liao, J.C. (2014). Afferent and motoneuron activity in response to single neuromast stimulation in the posterior lateral line of larval zebrafish. Journal of Neurophysiology. 112(6), 1329–39.

Hirono, M., Denis, C.S., Richardson, G.P. & Gillespie, P.G. (2004). Hair cells require phosphatidylinositol 4,5-bisphosphate for mechanical transduction and adaptation. Neuron. 44(2), 309–20.

Holt, J.R., Corey, D.P. & Eatock, R.A. (1997). Mechanoelectrical transduction and adaptation in hair cells of the mouse utricle, a low-frequency vestibular organ. The Journal of Neuroscience. 17(22), 8739–48.

Holt, J.R., Gillespie, S.K., Provance, D.W., Shah, K., Shokat, K.M., Corey, D.P., Mercer, J.A. & Gillespie, P.G. (2002). A chemical-genetic strategy implicates myosin-1c in adaptation by hair cells. Cell. 108(3), 371–81.

Holton, T. & Hudspeth, A.J. (1986). The transduction channel of hair cells from the bull-frog characterized by noise analysis. The Journal of Physiology. 375, 195–227.

Howard, J. & Hudspeth, A.J. (1987). Mechanical relaxation of the hair bundle mediates adaptation in mechanoelectrical transduction by the bullfrog’s saccular hair cell. Proceedings of the National Academy of Sciences of the United States of America. 84(9), 3064–8.

Hudspeth, A.J. (2014). Integrating the active process of hair cells with cochlear function. Nat Rev Neurosci. 15, 600–614.

Kastner, D.B. & Baccus, S.A. (2011). Coordinated dynamic encoding in the retina using opposing forms of plasticity. Nature Neuroscience. 14(10), 1317–22.

Kastner, D.B. & Baccus, S.A. (2013). Spatial segregation of adaptation and predictive sensitization in retinal ganglion cells. Neuron. 79(3), 541–54.

Keen, E.C. & Hudspeth, A.J. (2006). Transfer characteristics of the hair cell’s afferent synapse. Proceedings of the National Academy of Sciences of the United States of America. 103(14), 5537–42.

Kennedy, H.J., Evans, M.G., Crawford, A.C., and Fettiplace, R. (2003). Fast adaptation of mechanoelectrical transducer channels in mammalian cochlear hair cells. Nat Neuroscience. 6, 832–836.

Kindt, K.S., Finch, G. & Nicolson, T. (2012). Kinocilia mediate mechanosensitivity in developing zebrafish hair cells. Dev Cell. 23(2), 329–41.

Kros, C.J., Marcotti, W., van Netten, S.M., Self, T.J., Libby, R.T., Brown, S.D., Richardson, G.P. & Steel, K.P. (2002). Reduced climbing and increased slipping adaptation in cochlear hair cells of mice with Myo7a mutations. Nature Neuroscience. 5(1), 41–7.

Li, G.L., Keen, E., Andor-Ardó, D., Hudspeth, A.J. & von Gersdorff, H. (2009). The unitary event underlying multiquantal EPSCs at a hair cell’s ribbon synapse. The Journal of Neuroscience. 29(23), 7558–68.

Lister, J.A., Robertson, C.P., Lepage, T., Johnson, S.L. & Raible, D.W. (1999). nacre encodes a zebrafish microphthalmia-related protein that regulates neural-crest-derived pigment cell fate. Development. 126(17), 3757–67.

Lv, C., Stewart, W.J., Akanyeti, O., Frederick, C., Zhu, J., Santos-Sacchi, J., Sheets, L., Liao, J.C. & Zenisek, D. (2016). Synaptic Ribbons Require Ribeye for Electron Density, Proper Synaptic Localization, and Recruitment of Calcium Channels. Cell Rep. 15:2784–2795.

Maeda, R., Pacentine, I.V., Erickson, T. & Nicolson, T. (2017). Functional Analysis of the Transmembrane and Cytoplasmic Domains of Pcdh15a in Zebrafish Hair Cells. The Journal of Neuroscience. 37(12), 3231–45.

Marcotti, W. & Masetto, S. (2010). Hair Cells. Encyclopedia of Life Sciences.

Markin, V.S. & Hudspeth, A.J. (1995). Gating-spring models of mechanoelectrical transduction by hair cells of the internal ear. Annual Review of Biophysics and Biomolecular Structure. 24, 59–83.

Marvin, J.S., Borghuis, B.G., Tian, L., Cichon, J., Harnett, M.T., Akerboom, J., Gordus, A., Renninger, S.L., Chen, T.W., Bargmann, C.I., Orger, M.B., Schreiter, E.R., Demb, J.B., Gan, W.B., Hires, S.A. & Looger, L.L. (2013). An optimized fluorescent probe for visualizing glutamate neurotransmission. Nature Methods. 10(2), 162–70.

Masland, R.H. (2012). The neuronal organization of the retina. Neuron. 76(2), 266–80.

McHenry, M.J. & van Netten, S.M. (2007). The flexural stiffness of superficial neuromasts in the zebrafish (Danio rerio) lateral line. J Exp Biol. 210(Pt 23), 4244–53.

McHenry, M.J., Feitl, K.E., Strother, J.A. & Van Trump, W.J. (2009). Larval zebrafish rapidly sense the water flow of a predator’s strike. Biol Lett. 5(4), 477–9.

McHenry, M.J., Strother, J.A. & van Netten, S.M. (2008). Mechanical filtering by the boundary layer and fluid-structure interaction in the superficial neuromast of the fish lateral line system. J Comp Physiol A Neuroethol Sens Neural Behav Physiol. 194(9), 795–810.

Metcalfe, W.K., Kimmel, C.B. & Schabtach, E. (1985). Anatomy of the posterior lateral line system in young larvae of the zebrafish. Journal of Comparative Neurology. 233(3), 377–89.

Nicolson, T. (2012). The Hair Cell Synapse. in Synaptic Mechanisms in the Auditory System. pp. 43–60.

Nicolson, T. (2015). Ribbon synapses in zebrafish hair cells. Hear Res. 330, 170–177.

Nikolaev, A., Leung, K.M., Odermatt, B. & Lagnado, L. (2013). Synaptic mechanisms of adaptation and sensitization in the retina. Nature neuroscience. 16(7), 934–41.

Nouvian, R., Beutner, D., Parsons, T.D. & Moser, T. (2006). Structure and function of the hair cell ribbon synapse. J Membr Biol. 209(2-3), 153–65.

Odermatt, B., Nikolaev, A. & Lagnado, L. (2012). Encoding of luminance and contrast by linear and nonlinear synapses in the retina. Neuron. 73(4), 758–73.

Ohn, T.L., Rutherford, M.A., Jing, Z., Jung, S., Duque-Afonso, C.J., Hoch, G., Picher, M.M., Scharinger, A., Strenzke, N. & Moser, T. (2016). Hair cells use active zones with different voltage dependence of Ca2+ influx to decompose sounds into complementary neural codes. Proceedings of the National Academy of Sciences of the United States of America. 113(32), E4716–25.

Olive, R., Wolf, S., Dubreuil, A., Bormuth, V., Debrégeas, G. & Candelier, R. (2016). Rheotaxis of Larval Zebrafish: Behavioral Study of a Multi-Sensory Process. Frontiers in systems neuroscience. 10, p. 14.

Olt, J., Johnson, S.L. & Marcotti, W. (2014). In vivo and in vitro biophysical properties of hair cells from the lateral line and inner ear of developing and adult zebrafish. The Journal of Physiology. 592(Pt 10), 2041–58.

Olt, J., Ordoobadi, A.J., Marcotti, W. & Trapani, J.G. (2016). Physiological recordings from the zebrafish lateral line. Methods Cell Biol. 133, 253–79.

Oteiza, P., Odstrcil, I., Lauder, G., Portugues, R. & Engert, F. (2017). A novel mechanism for mechanosensory-based rheotaxis in larval zebrafish. Nature. 547, 445–448.

Palmer, L.M., Deffenbaugh, M. & Mensinger, A.F. (2005). Sensitivity of the anterior lateral line to natural stimuli in the oyster toadfish, Opsanus tau (Linnaeus). J Exp Biol. 208(Pt 18), 3441–50.

Pangršič, T., Gabrielaitis, M., Michanski, S., Schwaller, B., Wolf, F., Strenzke, N. & Moser, T. (2015). EF-hand protein Ca2+ buffers regulate Ca2+ influx and exocytosis in sensory hair cells. Proceedings of the National Academy of Sciences of the United States of America. 112(9), E1028–37.

Peng, A.W., Effertz, T. & Ricci, A.J. (2013). Adaptation of mammalian auditory hair cell mechanotransduction is independent of calcium entry. Neuron. 80(4), 960–72.

Platzer, J., Engel, J., Schrott-Fischer, A., Stephan, K., Bova, S., Chen, H., Zheng, H. & Striessnig, J. (2000). Congenital deafness and sinoatrial node dysfunction in mice lacking class D L-type Ca2+ channels. Cell. 102(1), 89–97.

Pujol-Martí, J. & López-Schier, H. (2013). Developmental and architectural principles of the lateral-line neural map. Front Neural Circuits. 7, p. 47.

Pujol-Martí, J., Zecca, A., Baudoin, J.P., Faucherre, A., Asakawa, K., Kawakami, K. & López-Schier, H. (2012). Neuronal birth order identifies a dimorphic sensorineural map. The Journal of Neuroscience. 32(9), 2976–87.

Ricci, A.J. & Kachar, B. (2007). Hair Cell Mechanotransduction: The Dynamic Interplay Between Structure and Function, in Current Topics in Membranes. Elsevier. 339–74.

Ricci, A.J., Bai, J.P., Song, L., Lv, C., Zenisek, D. & Santos-Sacchi, J. (2013). Patch-clamp recordings from lateral line neuromast hair cells of the living zebrafish. The Journal of Neuroscience. 33(7), 3131–4.

Russell, I.J. & Roberts, B.L. (1974). Active reduction of lateral-line sensitivity in swimming dogfish. Journal of Comparative Physiology A: Neuroethology, Sensory, Neural, and Behavioral Physiology. 94(1), 7–15.

Schnee, M.E., Lawton, D.M., Furness, D.N., Benke, T.A. & Ricci, A.J. (2005). Auditory hair cell-afferent fiber synapses are specialized to operate at their best frequencies. Neuron. 47(2), 243–54.

Schnee, M.E., Santos-Sacchi, J., Castellano-Muñoz, M., Kong, J.H. & Ricci, A.J. (2011). Calcium-dependent synaptic vesicle trafficking underlies indefatigable release at the hair cell afferent fiber synapse. Neuron. 70(2), 326–38.

Sheets, L., He, X.J., Olt, J., Schreck, M., Petralia, R.S., Wang, Y.X., Zhang, Q., Beirl, A., Nicolson, T., Marcotti, W., Trapani, J.G. & Kindt, K.S. (2017). Enlargement of Ribbons in Zebrafish Hair Cells Increases Calcium Currents But Disrupts Afferent Spontaneous Activity and Timing of Stimulus Onset,. The Journal of Neuroscience. 37(26), 6299–313.

Sheets, L.,Kindt, K.S. & Nicolson, T. (2012). Presynaptic CaV1.3 channels regulate synaptic ribbon size and are required for synaptic maintenance in sensory hair cells. The Journal of Neuroscience. 32(48), 17273–86.

Shepherd, G.M. & Corey, D.P. (1994). The extent of adaptation in bullfrog saccular hair cells. The Journal of Neuroscience. 14(10), 6217–29.

Stauffer, E.A. & Holt, J.R. (2007). Sensory transduction and adaptation in inner and outer hair cells of the mouse auditory system. Journal of neurophysiology. 98(6), 3360–9.

Stauffer, E.A., Scarborough, J.D., Hirono, M., Miller, E.D., Shah, K., Mercer, J.A., Holt, J.R. & Gillespie, P.G. (2005). Fast adaptation in vestibular hair cells requires myosin-1c activity. Neuron. 47(4), 541–53.

Stewart, W.J., Cardenas, G.S. & McHenry, M.J. (2013). Zebrafish larvae evade predators by sensing water flow. J Exp Biol. 216(Pt 3), 388–98.

Striessnig, J., Bolz, H.J. & Koschak, A. (2010). Channelopathies in Cav1.1, Cav1.3, and Cav1.4 voltage-gated L-type Ca2+ channels. Pflugers Archiv: European journal of physiology. 460(2), 361–74.

Suli, A., Watson, G.M., Rubel, E.W. & Raible, D.W. (2012). Rheotaxis in larval zebrafish is mediated by lateral line mechanosensory hair cells. PloS One. 7(2), p. e29727.

Trapani, J.G. & Nicolson, T. (2011). Mechanism of spontaneous activity in afferent neurons of the zebrafish lateral-line organ. The Journal of Neuroscience. 31(5), 1614–23.

Trapani, J.G., Obholzer, N., Mo, W., Brockerhoff, S.E. & Nicolson, T. (2009). Synaptojanin1 is required for temporal fidelity of synaptic transmission in hair cells. PLoS Genet. 5(5), p. e1000480.

Troconis, E.L., Ordoobadi, A.J., Sommers, T.F., Aziz-Bose, R., Carter, A.R. & Trapani, J.G. (2017). Intensity-dependent timing and precision of startle response latency in larval zebrafish. The Journal of Physiology. 595(1), 265–82.

Voigt, R., Carton, A.G. & Montgomery, J.C. (2000). Responses of anterior lateral line afferent neurones to water flow. J Exp Biol. 203(Pt 16), 2495–502.

Vollrath, M.A. & Eatock, R.A. (2003). Time course and extent of mechanotransducer adaptation in mouse utricular hair cells: comparison with frog saccular hair cells. Journal of neurophysiology. 90(4), 2676–89.

Wark, B., Lundstrom, B.N. & Fairhall, A. (2007). Sensory adaptation, Current opinion in neurobiology. 17(4), 423–9.

Weisz, C.J., Lehar, M., Hiel, H., Glowatzki, E. & Fuchs, P.A. (2012). Synaptic transfer from outer hair cells to type II afferent fibers in the rat cochlea. The Journal of Neuroscience. 32(28), 9528–36.

Zhang, Q.X., He, X.J., Wong, H.C. & Kindt, K.S. (2016). Functional calcium imaging in zebrafish lateral-line hair cells. Methods Cell Biol. 133, 229–52.

